# PI3K inhibition as a novel therapeutic strategy for neoadjuvant chemoradiotherapy resistant oesophageal adenocarcinoma

**DOI:** 10.1101/2020.10.23.351981

**Authors:** Sarah D. Edge, Isaline Renard, Emily Pyne, Hannah Moody, Rajarshi Roy, Andrew W. Beavis, Stephen J. Archibald, Christopher J. Cawthorne, Stephen G. Maher, Isabel M. Pires

**Affiliations:** Hypoxia and Tumour Microenvironment Lab, Department of Biomedical Sciences, Faculty of Health Sciences, University of Hull, UK; Positron Emission Tomography Centre, Department of Biomedical Sciences, Faculty of Health Sciences, University of Hull, UK; Institute of Cancer Therapeutics; School of Medicine and Medical Sciences, University of Bradford, UK; Queen’s Centre for Oncology and Haematology, Castle Hill Hospital, Castle Rd, Cottingham HU16 5JQ, UK; Department of Medical Physics, Queen’s Centre for Oncology, Hull University Teaching Hospitals NHS Trust, Cottingham, HU16 5JQ, UK; Faculty of Health Sciences, University of Hull, Cottingham road, Hull, HU16 7RX, UK; Faculty of Health and Well Being, Sheffield-Hallam University, Collegiate Crescent, Sheffield, S10 2BP; Department of Surgery, Trinity Translational Medicine Institute, Trinity College Dublin, St. James’s Hospital, Dublin, Ireland; Nuclear Medicine and Molecular Imaging, Department of Imaging and Pathology, KU Leuven, 3000 Leuven, Belgium

**Keywords:** oesophageal cancer, PI3K, miR-187, radiation, PTEN

## Abstract

**Objectives:** Neoadjuvant chemo-radiotherapy (neo-CRT) prior to surgery is the standard of care for oesophageal adenocarcinoma (OAC) patients. Unfortunately, most patients fail to respond to treatment. MiR-187 was previously shown to be downregulated in neo-CRT non-responders, whist *in vitro* miR-187 overexpression enhanced radio-sensitivity and upregulated *PTEN*. This study evaluates the role of miR-187 and downstream PI3K signalling in radiation response in OAC.

**Methods:** The effect of miR-187 overexpression on downstream PI3K signalling was evaluated in OAC cell lines by qPCR and western blotting. *PTEN* expression was analysed in OAC pre-treatment biopsies of neo-CRT responders and non-responders. Pharmacological inhibition of PI3K using GDC-0941 was evaluated in combination with radiotherapy in 2D and 3D OAC models *in vitro* and as a single agent *in vivo*. Radiation response *in vitro* was assessed via clonogenic assay.

**Results:** *PTEN* expression was significantly decreased in neo-CRT non-responders. MiR-187 overexpression significantly upregulated *PTEN* expression and inhibited downstream PI3K signalling *in vitro*. GDC-0941 significantly reduced viability and enhanced radiation response *in vitro* and led to tumour growth inhibition as a single agent *in vivo*.

**Conclusions:** Targeting of PI3K signalling is a promising therapeutic strategy for OAC patients who have repressed miR-187 expression and do not respond to conventional neo-CRT.

**Advances in knowledge:** This is the first study evaluating the effect of PI3K inhibition on radio-sensitivity in OAC, with a particular focus on patients that do not respond to neo-CRT. We have shown for the 1^st^ time that targeting of PI3K signalling is a promising alternative therapeutic strategy for OAC patients who do not respond to conventional neo-CRT.

## Introduction

Oesophageal cancer has extremely poor prognosis and is the seventh most common and sixth most lethal cancer worldwide (1). Oesophageal adenocarcinoma (OAC) is the most prevalent histological subtype in western countries, with rapidly rising incidence rates (2),(3). Neoadjuvant chemoradiotherapy (neo-CRT) is becoming the standard of care for patients with locally advanced OAC (4). Patients achieving a pathological complete response (pCR) upon neo-CRT have been shown to have five-year survival rates up to 50% (5). Unfortunately, less than one third of OAC patients treated with neo-CRT achieve pCR (6). Therefore, identification of predictive therapy response biomarkers and novel therapeutic strategies is essential to improve patient prognosis.

MicroRNAs (miRNAs) are gene expression regulators, identified as predictors and modulators of treatment response in OAC (7). We have previously demonstrated that miR-31, miR-330-5p and miR-187 are significantly downregulated in pre-treatment tumour biopsies from OAC patients who are poor responders to neo-CRT (8–10). MiR-187 overexpression significantly enhanced sensitivity to radiotherapy and cisplatin *in vitro* and altered gene expression patterns, including *PTEN* (10). *PTEN* is a negative regulator of the phosphatidylinositol 3 kinase (PI3K)-protein kinase B (AKT) signalling pathway, whose hyperactivation is implicated cancer development and progression (11). Activation of PI3K-AKT signalling has been demonstrated in OAC (12), although the potential clinical relevance of PI3K inhibitors remains poorly understood.

GDC-0941 is a potent pan-PI3K inhibitor (13). Pre-clinical studies and phase I clinical trials have demonstrated that GDC-0941 promotes anti-tumorigenic effects and is suitable for treatment of solid tumours (13–16). GDC-0941 was also shown to enhance radio-sensitivity in glioblastoma multiforme *in vitro* and in thyroid carcinoma *in vivo* (17, 18). However, anti-tumorigenic properties of GDC-0941 have not previously been assessed in OAC.

In this study, we investigated the role of miR-187 and the PI3K axis in modulating response to neo-CRT. We showed that *PTEN* expression is downregulated in OAC patients who are poor responders to neo-CRT, and that miR-187 overexpression increases *PTEN* expression and downregulates p-AKT levels in OAC cells. We then targeted PI3K signalling in OAC cell lines using GDC-0941, which led to decreased cell survival, increased radio-sensitivity *in vitro,* and inhibition of *in vivo* tumour growth for the first time in OAC. Taken together, we are demonstrating for the 1^st^ time that targeting PI3K in OAC is a promising therapeutic strategy in OAC and could improve outcome of patients who have repressed miR-187 expression and do not respond to conventional neo-CRT.

## Methods and Materials

### Cell culture, transfections, drug treatments, and irradiation treatment

OAC OE33, OE19 and SK-GT-4 cell lines were purchased from ECACC (UK). Cells were incubated at 37°C, 95% humidified air and 5% CO2. Spheroids were generated by seeding 2.5×10^4^ cells per well in ultra-low adherence round-bottomed 96-well plates (Corning). Media was replaced every 2 days. Spheroids were imaged using the GelCount instrument (Oxford Optronix), and spheroid size determined as previously reported (19, 20).

Ambion pre-miR miRNA precursor molecules (Thermo Fisher Scientific, UK) were used for transient miR-187 overexpression. GDC-0941 stock solutions (Abcam, UK; Selleckchem, UK) were prepared in DMSO (dimethyl sulfoxide). X-ray irradiation was carried out using a RS-2000 biological irradiator (Rad Source Technologies, Georgia, USA) at a 1.87 Gy/min dose rate, calibrated to National Physics Laboratory (NPL) standards (21).

### Patient treatment, tissue collection, and histology

Following ethical approval (Joint St James’s Hospital/AMNCH ethical review board, Reference ID 2011/27/01) and written informed consent, diagnostic biopsy tumour specimens were taken from patients with a diagnosis of operable OAC, prior to neo-CRT. All patients received a complete course of neo-CRT, consisting of two courses of 5-fluorouracil (5-FU) and cisplatin, plus 40.05 Gy in 15 daily fractions (2.67 Gy/fraction) over 3 weeks as previously described (22). Diagnostic endoscopic biopsies were obtained prior to neo-CRT. Immediately adjacent tissue was taken for histologic confirmation, which was performed using routine Haematoxylin and Eosin staining. Specimens were immediately placed in RNAlater (Ambion) and refrigerated for 24 h, before removal of RNA later and storage at −80°C.

All specimens were assessed by an experienced pathologist. Tumour response to treatment was assigned 1 of 5 tumour regression grades (TRG) as previously described (23). Good responders were classified as achieving a TRG of 1 or 2, whilst poor responders were classified as having a TRG of 3, 4 or 5, as previously described (9). For the purposes of this study patients with a TRG 3 were excluded.

### *In vivo* studies

Xenograft models were generated through the subcutaneous implant of 5×10^6^ OE33 cells into the right-hand side of the middorsal region of the back of 16 x 52-58 day old female CB17.Cg-Prkdc^scid^Lyst^bg-J^ SCID-Bg mice (sourced from Charles River UK). Animals were anaesthetised using 2% isoflurane, and a small area of fur shaved around the implant site prior to implant. The cell suspension was prepared in a 1:1 ratio of serum free RPMI medium and Cultrex BME (24), prior to implant of 0.1 mL of the 5×10^7^ cells/ mL suspension. Tumour measurements were taken three times weekly with callipers and animals randomly assigned to control and treatment groups when tumour volume reached approximately 300-350mm^3^ (14, 25, 26). Three animals developed tumours at a later date and were not included in the analysis of tumour growth. Twice daily treatments of GDC-0941 or vehicle control (0.1 mL/ 10g) were administered via oral gavage at a concentration of 50 mg/kg for 8 days, as previously reported {Burrows, 2011 #23;Burrows, 2013 #32;Cawthorne, 2013 #33. Health status and weight of each animal were monitored throughout the experiment. Animals were sacrificed via a schedule 1 method (dislocation of the neck), and tumours were excised and snap frozen. All procedures were carried out in accordance with the Animals (Scientific Procedures) Act 1986 and UKCCCR Guidelines 2010 using protocols approved by the University of Hull Animal Welfare and Ethical Review Body (AWERB) under Home Office Project Licence 60/4549. The study was powered using effect sizes and tumour growth variability seen in previous experiments {Burrows, 2011 #23;Burrows, 2013 #32;Cawthorne, 2013 #33 using G*Power software (27). Tumour growth over the entire course of treatment was compared between groups.

### RT-qPCR

Xenograft tumour tissue samples and patient tumour biopsies RNA was isolated using an Allin-One purification kit (Norgen Biotek). Aurum Total RNA Mini Kit (Biorad, UK) was used for RNA extraction from cell lines.

For miRNA expression analysis, 10 ng total RNA was first reverse transcribed to cDNA using the TaqMan MicroRNA Reverse Transcription Kit (Applied Biosystems, UK). TaqMan microRNA assays and TaqMan 2X Universal PCR Master Mix (Applied Biosystems, UK) were used, with RNU48 used as the endogenous control. For mRNA analysis, 1μg total RNA was reversed transcribed using the RevertAid H Minus First Strand cDNA Synthesis Kit (Thermo Scientific, UK). QuantiNova SYBR Green PCR Kit and QuantiTECT primer assays (Qiagen, UK) were used, with *B2M* or *18S* used as the endogenous control. RT-qPCR was performed using the Step One Plus Real Time PCR System (Applied Bioscience, USA) and data was analysed using the 2^-ΔΔCt^ method (28). Primer details available in Supplementary Table 1.

### Clonogenic assay

Cells were plated at a density of 500-5000 cells/well in 6-well plastic plates. Colonies were fixed and stained by crystal violet staining solution (0.1% w/v crystal violet, 70% v/v methanol, 30% v/v dH_2_O) after 7-10 days post treatment. Colonies were counted using a Gel Count system (Oxford Optronics, UK).

### Cell viability assay

Short term cell viability was assessed using the Cell Titer 96 AQeous One Solution MTS cell proliferation assay (Promega, UK), as previously described {Beeby, 2020 #46}.

### Immunoblotting

Cells lysates were prepared in UTB as previously reported (29). Tumour samples were prepared for lysis using a BioPulverizer and Cryo-cup grinder (BioSpec, USA) and lysed in RIPA buffer (Cell Signalling Technology, USA) supplemented with protease (Mini, EDTA-free Protease Inhibitor Cocktail, Roche, UK) and phosphatase (PhosSTOP, Roche, UK) inhibitors. Western blotting was performed as previously described (29). Densitometric analysis of band intensity of blots was carried out using Image J (NIH, USA).

### Statistical analysis

Statistical analysis was carried out using the GraphPad PRISM (GraphPad software Inc, California, USA) and the data presented represent the mean +/- the standard error of the mean (SEM). Statistical analysis was carried out using an unpaired Student’s T-test or a 2-way ANOVA. Significance was assumed if p<0.05.

**Detailed materials and methods are described in the Supplementary Materials.**

## Results

### PTEN expression is associated with CRT response in OAC patients

We have previously shown that miR-187 is downregulated in poor responder patients in OAC and can impact on gene expression for survival pathways (10). Here, we confirmed that miR-187 overexpression resulted in significant upregulation of *PTEN* mRNA expression in a OAC cell line panel compared to non-transfected controls (Figure 1A, Supplementary Figure 1A), linking miR-187 levels with *PTEN* expression. As a search of the miR-code database (30) did not reveal a miR-187 binding site in the *PTEN* promotor, *PTEN* regulatory co-factors were identified using the UCSC genome browser, and those with miR-187 seed sites identified using the miR-code database (Supplementary Table 3). For the most promising negative regulators *EP300* or *DNMT1*, expression was not affected by miR-187 overexpression, while *KDMB5* expression was significantly increased in the OE33 and SK-GT-4 cell lines (Supplementary Figure 2), indicating that other factors must be responsible for miR-187-mediated *PTEN* modulation. The effect of miR-187 overexpression on downstream PI3K-AKT signalling was then assessed through analysis of phosphorylated AKT levels (Serine 473). MiR-187 overexpression resulted in significant reduction of pAKT levels compared to the nontransfected controls (Figure 1B, Supplementary Figure 3). Finally, we observed *PTEN* expression was significantly downregulated in the pre-treatment tumour biopsies of poor responders to neo-CRT compared to the responder group (Figure 1C). These data suggest that PTEN/PI3K-AKT signalling correlates with miR-187 expression *in vitro* and is associated with radiation response efficacy in OAC patients.

**Figure 1.**
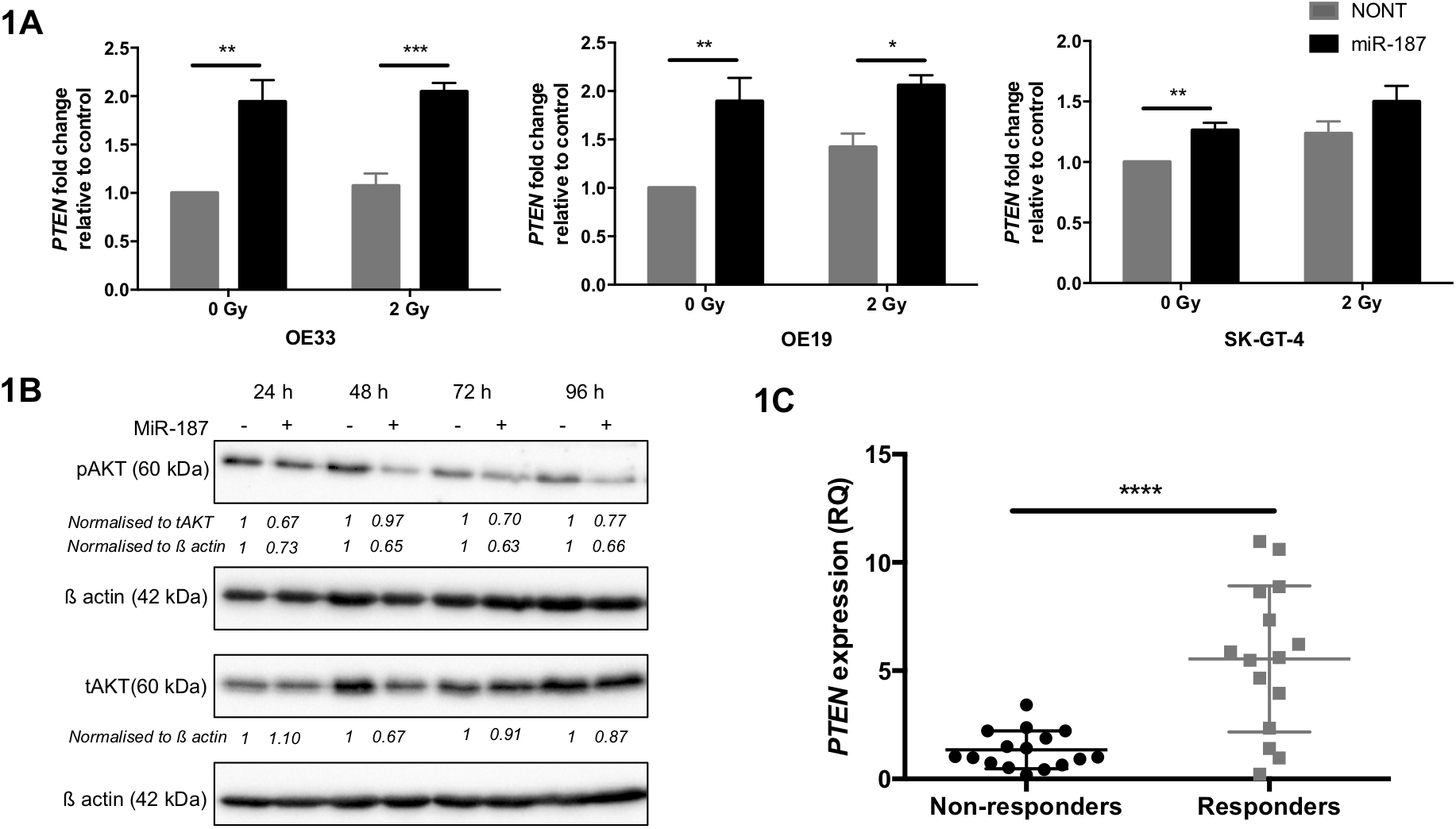
PTEN expression is associated with CRT response in OAC patients. A) OE33, OE19 and SK-GT-4 cells were transfected with pre-miR-187 precursor molecules and exposed to 2 Gy irradiation. PTEN fold change was analyzed by qPCR and expression calculated relative to the nontransfected control (NONT). Histogram represents an average of n=3 experimental repeats. B) OE33 cells were transfected with pre-miR-187 precursor molecules and tAKT/ pAKT protein levels assessed by western blot. Western blot and densitometry shown is representative of n=3 repeats. C) PTEN expression was assessed in the pre-treatment diagnostic biopsy specimens of OAC patients (n=3l) by qPCR. Patients were classified as neo-CRT responders (TRG 1 and 2, n=l5) and non-responders (TRG 4 and 5, n=l6). Error bars represent mean ± SEM. *p < 0.05, **p< 0.01, ***p<0.001, ****p<0.0001.

### PI3K inhibition efficiently reduces OAC cell survival *in vitro* and *in vivo* tumour growth

We hypothesised that pharmacological inhibition of PI3K may provide an alternative novel therapeutic strategy for OAC in patients with repressed miR-187 expression who do not respond to neo-CRT. Clinically relevant PI3K inhibitor GDC-0941 was selected for further investigation due to the high efficacy shown in pre-clinical studies and phase I clinical trials for solid tumours (13–15, 25, 26, 31). GDC-0941 treatment resulted in PI3K inhibition as indicated via a reduction (lower doses) or abrogation (higher doses) in phospho-AKT in the OAC cell line panel (Figure 2A). Treatment with GDC-0941 significantly reduced both shortterm viability(Figure 2B) and long-term clonogenic survival (Figure 2C) for the OAC cell line panel. These data confirm that GDC-0941 effectively reduces cell survival as a single agent *in vitro*.

**Figure 2.**
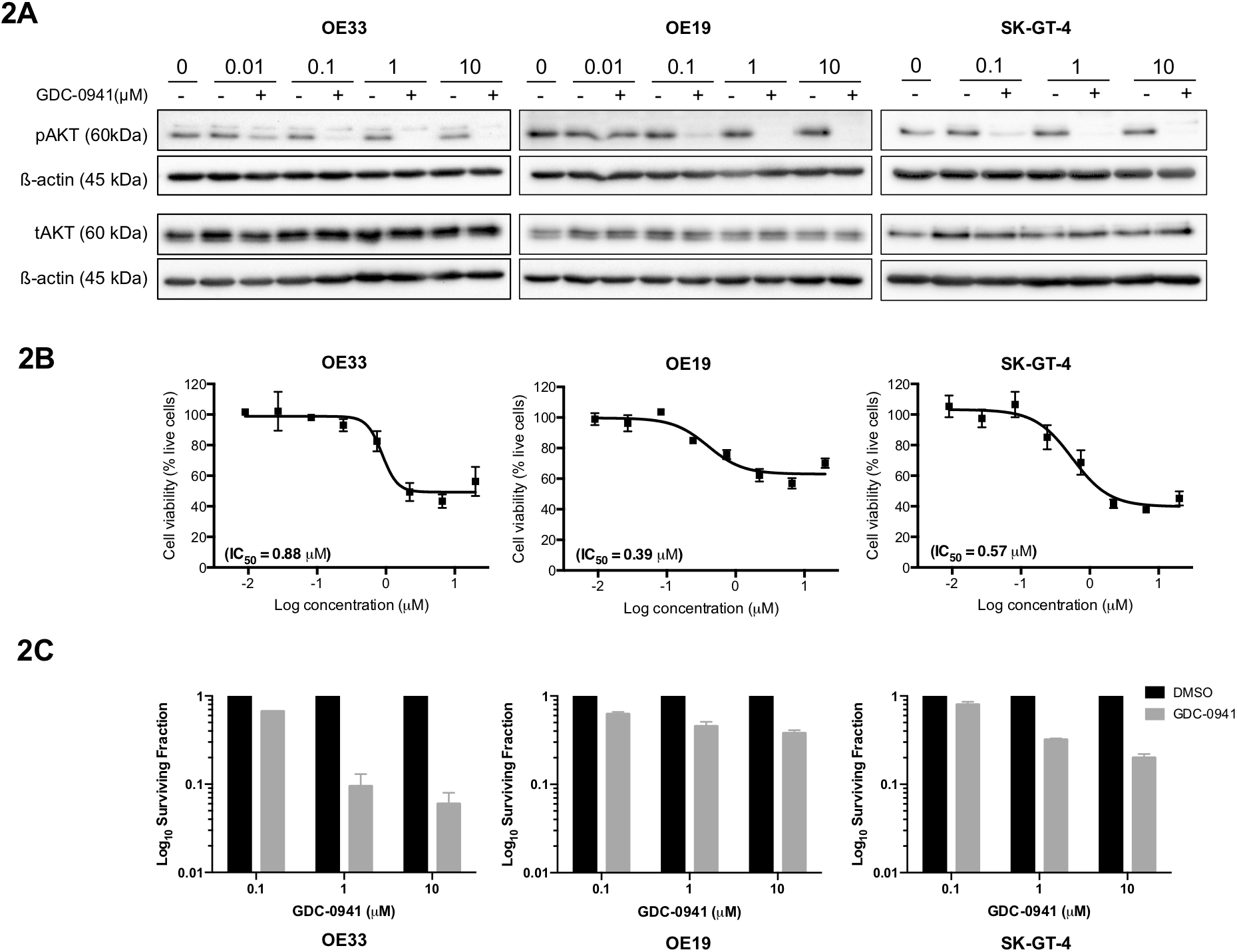
PI3K inhibition significantly reduces OAC proliferation and survival *in vitro*. A) Cells were treated with GDC-0941 or vehicle control for 18 h and tAKT/ pAKT (Ser 473) protein levels assessed by western blot. Western blot is representative of n=3 repeats. B) Cells were treated with GDC-0941 at a top concentration of 20 μM and an 8-point dose response curve generated. The MTS reagent was added to the plates 48 h post treatment and absorbance measured at 490 nm 4 h later. Cell viability was calculated relative to the corresponding vehicle control for each dose (n=3). C) Cells were treated with GDC-0941 or vehicle control and the effect on long term survival assessed using the clonogenic assay. Data represent the average of n=2 experimental repeats. Error bars represent mean ± SEM.

Pre-clinical efficacy was then assessed *in vivo* using OE33 tumour xenograft models. Significant reduction in tumour growth was observed in mice treated with GDC-0941 (50 mg/kg) compared to those treated with the vehicle control (Figure 3A). pAKT levels were reduced in the GDC-0941 treated group compared to those treated with the vehicle control, indicating that PI3K signalling was successfully inhibited (Figure 3B). This is the first pre-clinical study to evaluate its efficacy for the treatment of OAC.

**Figure 3.**
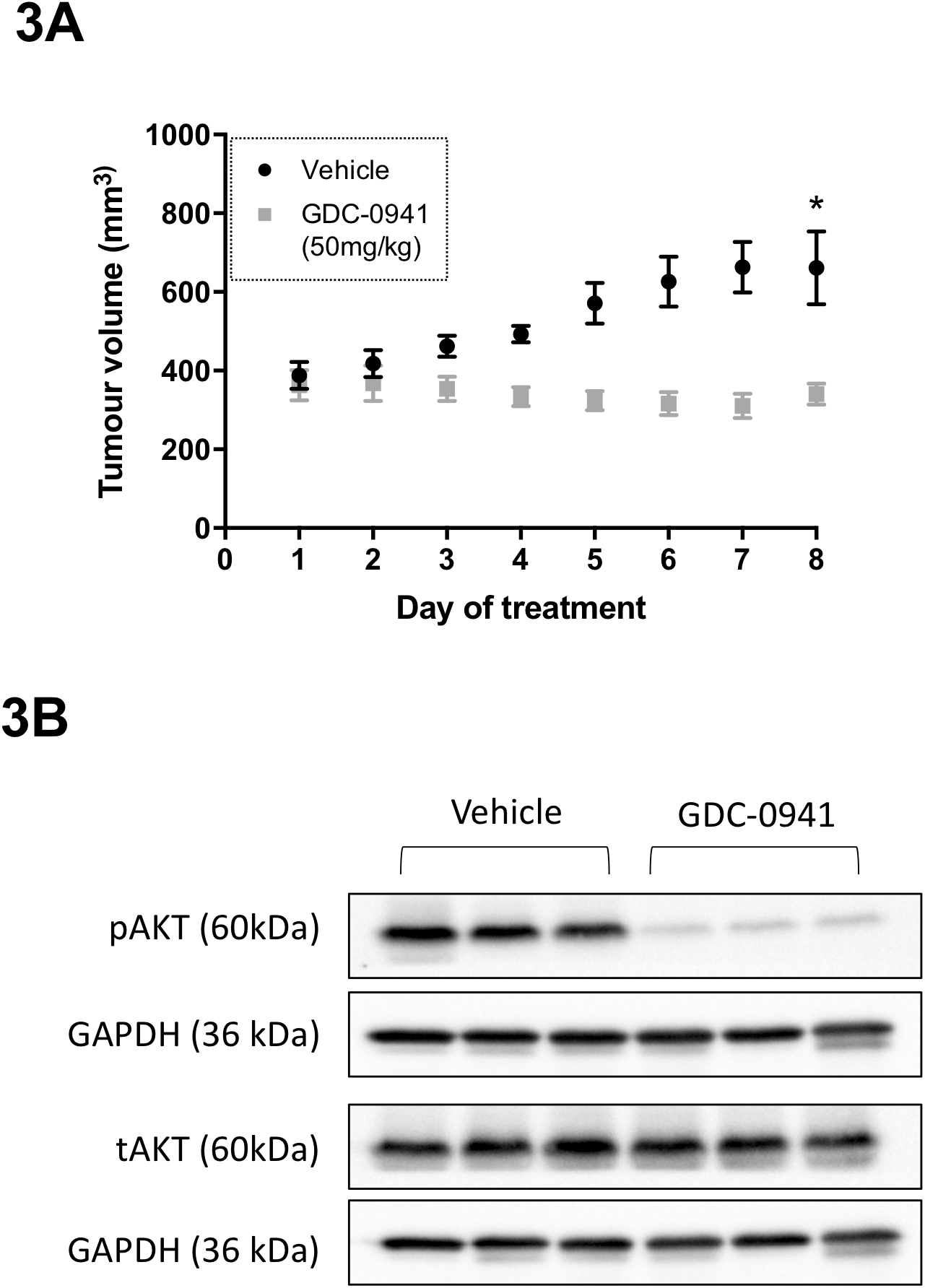
GDC-0941 treatment significantly decreases OAC tumour growth *in vivo*. A) SCID-Beige female mice bearing OE33 tumour xenografts were treated twice daily with GDC-0941 (50 mg/kg) or vehicle control via oral gavage for 8 days. Data represents the average mean tumour volume of n=3 (treated) or n=3 (vehicle) animals per treatment group. B) Animals were sacrificed 1 hour after the final treatment dose and tAKT/ pAKT protein levels assessed by western blot. Western blot is representative of n=3 repeats. Error bars represent mean ± SEM. *p < 0.05

### PI3K inhibition enhances radiosensitivity in OAC *in vitro*

Treatment with GDC-0941 significantly reduced long term clonogenic survival of OAC cell lines both as a single agent and in combination with irradiation as compared to the vehicle control, through an additive effect (Figure 4A, Supplementary Figure 4). Finally, treatment of all three OAC cell lines with GDC-0941 significantly reduced OAC spheroid volume, which was exacerbated in combination with ionising radiation (Figure 4B-D). These data suggest that

**Figure 4.**
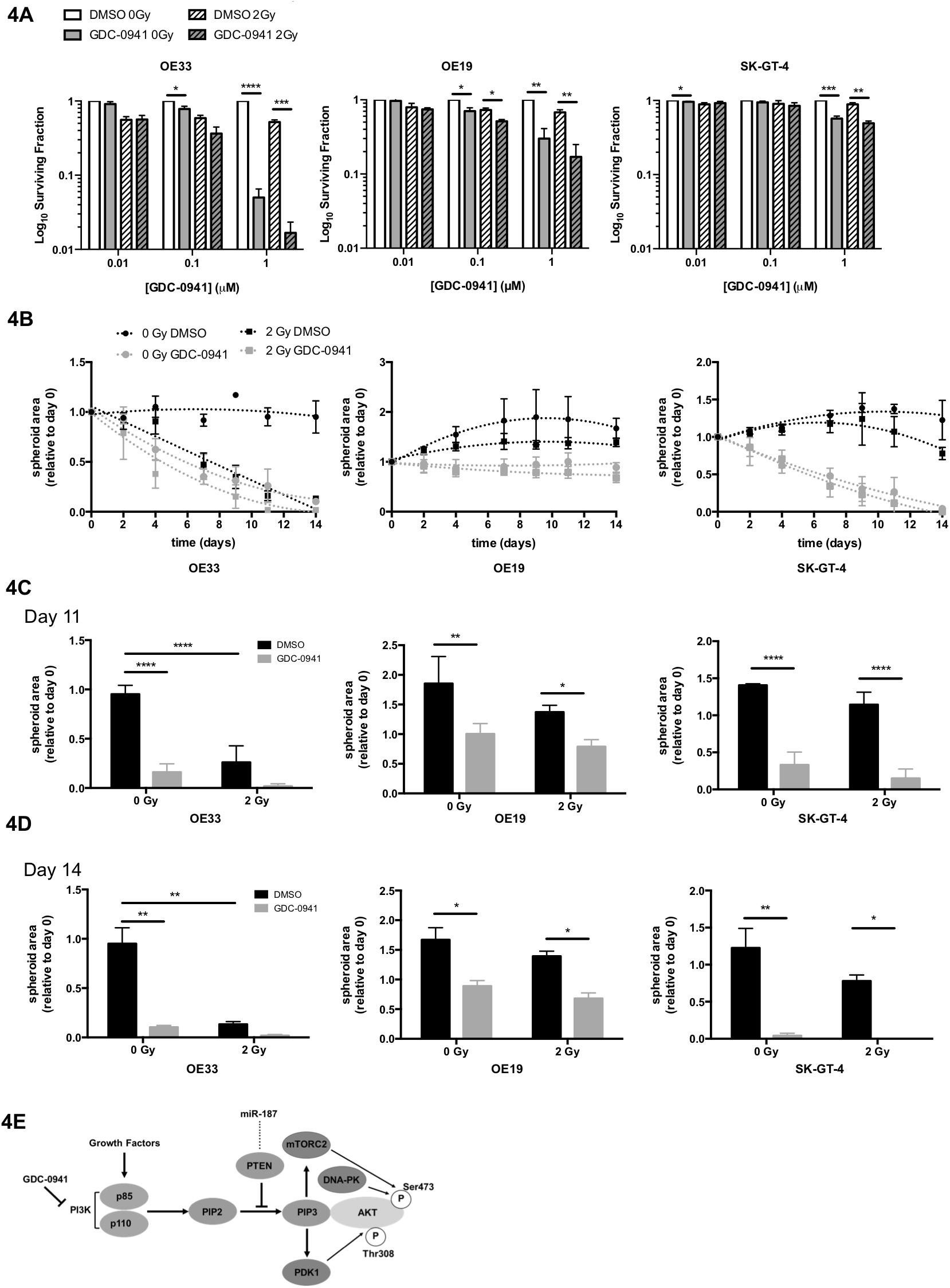
PI3K inhibition enhances radiotherapy treatment in 2D and 3D OAC models. A) Cells were treated with GDC-0941 or vehicle control for 18 hours and exposed to 2 Gy irradiation. The effect of GDC-0941 as a single agent and in combination with 2 Gy irradiation was assessed using the clonogenic assay. The surviving fraction was calculated relative to the 0 Gy DMSO control. B-D) OE33, OE19, and SK-GT-4 spheroids were treated with 1 μM GDC-0941 or DMSO for 21 days in combination with a total of 5×2 Gy irradiation fractions or mock irradiated for the first 5 days. Spheroids were imaged using the GelCount instrument (Oxford Optronix), and spheroid size determined using ImageJ (NIH). Spheroid area is shown relative to day 0. Error bars represent mean ± SEM. *p < 0.05, **p< 0.01, ***p<0.001, ****p<0.0001. E) Schematic of the effect of GDC-0941 treatment of the PI3K-AKT signalling pathway.

PI3K inhibition through treatment with GDC-0941 significantly reduces the long-term clonogenic survival of OAC cell lines as a single agent and in combination with physiologically relevant levels of ionising radiation.

## Discussion

In the present study, miR-187 overexpression was shown to induce upregulation of the tumour suppressor gene *PTEN* which resulted in subsequent inhibition of downstream PI3K-AKT signalling in OAC cell lines. *PTEN* expression levels were also found to be significantly downregulated in the pre-treatment biopsy specimens of OAC patients displaying a poor response to neo-CRT (Figure 1). Our findings are supported by the findings of Saeed and colleagues, where AKT expression levels were increased in the pre-treatment tumour biopsies of OAC patients identified as partial or non-responders to neo-CRT compared to those with a pCR. AKT expression was also shown to significantly correlate to the degree of pathologic response (32). As *PTEN* is a well-established negative regulator of the PI3K-AKT signalling pathway, our current and previous findings, alongside those published by Saeed and colleagues (32), indicate that miR-187, *PTEN*, and downstream PI3K-AKT signalling are implicated in modulating neo-CRT response in OAC. This suggests that decreased PTEN expression could be predictive of OAC radioresistance, as it is well established that the PI3K-AKT signalling pathway becomes upregulated in response to radiotherapy and is implicated in radiation resistance (33). Therefore, manipulation of PI3K may provide alternative therapeutic strategies for tackling radiation resistance in OAC.

We hypothesised that a negative *PTEN* transcriptional regulator harbouring a miR-187 binding site is responsible for miR-187 modulation of *PTEN* in OAC, as PTEN does not appear to be a direct target for miR-187. Unfortunately, candidate regulators *EP300, KDM5B* and *DMNT1* were also not affected by miR-187 overexpression (Supplementary Figure 2), indicating that other intermediate factors might be involved. Of particular interest from our *in silico* analysis (Supplementary Table 3), both *NFIC* (Nuclear Factor I C) and *PHF8* (PHD Finger Protein 8) were identified as potential miR-187 targets with an impact on PTEN expression through a yet unknown regulation mechanism, and have been shown to be overexpressed in oesophageal squamous cell carcinoma (34, 35), which will be evaluated in future studies.

Inhibition of PI3K signalling using the clinically relevant agent GDC-0941 significantly decreased cellular survival and enhanced radiation response in 2D and 3D models *in vitro*, as well as leading to tumour growth inhibition as a single agent in OAC xenograft models (Figures 2–4). The data presented here support previous studies which demonstrated that OAC and OSCC cell lines are sensitive to the previous generation PI3K inhibitors Wortmannin and LY294002 (36–38). Unlike the previous generation PI3K inhibitors, the PI3K inhibitor GDC-0941 used in this study is a highly selective inhibitor that targets all four isoforms of class I PI3Ks, the effects of which have not previously been assessed in oesophageal cancer (13). Importantly, this is the first study to demonstrate the efficacy of GDC-0941 for the treatment of OAC, either as a single agent or in combination with radiotherapy. These findings support those of previous pre-clinical studies and clinical trials indicating that GDC-0941 is suitable for the treatment of solid tumours (13, 15, 16). PI3K inhibition has been shown to modulate radio-sensitivity in a range of malignancies including lung, prostate and thyroid carcinoma (39, 40), although this is the first study evaluating the effect of PI3K inhibition on radio-sensitivity in OAC. Our study indicates, for the first time, that PI3K inhibition may enhance radiotherapy response in OAC.

## Conclusions

We have shown that PTEN and PI3K signalling are implicated in modulating tumour response to neo-CRT in OAC patients. We have also demonstrated that the clinically relevant PI3K inhibitor GDC-0941 significantly decreased cellular survival and enhanced the efficacy of radiation treatment in OAC cell lines *in vitro*, as well as leading to inhibiting tumour growth as a single agent in *in vivo* xenograft models. Importantly, the preliminary data presented here demonstrate that PI3K inhibitors such as GDC-0941 may provide an alternative therapeutic strategy for the treatment of OAC, either as a single agent or in combination with neo-CRT.

## Acknowledgements

The authors would like to thank Ellie Beeby and Dr Becky Bibby for technical support.

