## Supplementary information for "PI3K inhibition as a novel therapeutic strategy for neoadjuvant chemoradiotherapy resistant oesophageal adenocarcinoma"

### Supplementary methods

Cell culture and spheroid generation and treatment

The human OAC cell lines OE33, OE19 and SK-GT-4 were purchased from the European Collection of Authenticated Cell Cultures, UK (ECACC). Cell lines were cultured in Roswell Park Memorial Institute 1640 (RPMI 1640) medium with L-glutamine (Gibco, Life Technologies, UK) supplemented with 10% (v/v) heat inactivated foetal bovine serum (FBS; Gibco, Life Technologies, UK). Cells were incubated at 37°C, with 95% humidified air and 5% $CO_2$. All cell lines tested negative for mycoplasma infection.

Spheroids were generated by seeding cells at a density of $2.5x10^4$ cells per well in ultra-low adherence round-bottomed 96-well plates (Corning). After aggregation, spheroids were treated according to specific experimental conditions, with media replaced every 2 days. Spheroids were imaged using the GelCount instrument (Oxford Optronix), and spheroid size was determined as previously reported [1,2]

Transfection with Pre-miR miRNA pre-cursor molecules

Ambion pre-miR miRNA precursor molecules (Thermo Fisher Scientific, UK) were used to achieve transient miR-187 overexpression. Cells were seeded into 6 cm dishes at a density of $0.8x10^6$ cells/ dish prior to reverse transfection with 5 nM Pre-miR-187 precursor molecules in combination with Lipofectamine RNAiMAX (Invitrogen, UK). Non-transfected control cells were concurrently seeded and treated using an identical protocol whereby the miRNA precursor molecules were substituted for the vehicle control (nuclease free water).

Patient treatment and histology

Following ethical approval (Joint St James's Hospital/AMNCH ethical review board, Reference ID 2011/27/01) and written informed consent, diagnostic biopsy tumour specimens were taken from patients with a diagnosis of operable OAC, prior to neoadjuvant therapy. All patients received a complete course of neoadjuvant CRT. Chemotherapy consisted of 2 courses of 5-fluorouracil (5-FU) and cisplatin, as previously described [3]. Patients received 40.05 Gy in 15 daily fractions (2.67 Gy/fraction) over 3 weeks as previously described [3]. Surgical resection was performed approximately one month following completion of the CRT regimen. All resected oesophagectomy specimens were assessed by an experienced pathologist. Tumour response to treatment was assigned 1 of 5 tumour regression grades (TRG) as previously described [4]. Good responders were classified as patients achieving a TRG of 1 or 2, whilst poor responders were classified as patients having a TRG of 3, 4 or 5, as previously described [5]. For the purposes of this study patients with a TRG 3 were excluded.

Tissue collection

Diagnostic endoscopic biopsies were obtained by a qualified endoscopist prior to neoadjuvant therapy. Immediately adjacent tissue was taken for histologic confirmation, which was performed using routine Haematoxylin and Eosin staining. Specimens were immediately placed in RNAlater (Ambion) and refrigerated for 24 h, before removal of RNA later and storage at -80$^{o}$C.

Clonogenic assay

Cell seeding densities were optimised for each treatment condition to ensure that at least 100 colonies, each consisting of at least 50 cells were counted. Cells were seeded directly into 6 well plates at the optimised seeding densities (500-5000 cells/well). Once adhered, cells were treated with GDC-0941 or the DMSO 18 hours and then irradiated/ mock irradiated. Clonogenic plates were then placed in the incubator 7-10 days, fixed, and stained through the application of crystal violet staining solution (0.1% w/v crystal violet, 70% v/v methanol, 30% v/v $dH_2O$). Colonies were counted using a Gel Count system (Oxford Optronics, UK). Plating efficiencies were calculated as follows: average colony number / number of cells seeded. The surviving fraction (SF) for each treatment group was then determined as follows: (average colony number/plating efficiency of control) X seeding density [6].

Cell viability assay

Short term cell viability was assessed using the Cell Titer 96 AQeous One Solution MTS cell proliferation assay (Promega, UK) as previously described [2]. Cells were seeded into 96 well plates at a seeding density of 3000 cells/well and incubated over night to allow cells to adhere. Complete medium was replaced with treatment conditions and plates were returned to the incubator for a further 48 h, and MTS reagent added as per manufacturer's instructions. Media only absorbance was subtracted from mean absorbance values and cell viability for each treatment condition was calculated relative to DMSO vehicle control, normalised to 100% viability.

<u>Western blotting</u>

Cells were lysed in UTB (9M Urea, 75 mM Tris- HCL pH 7.5, 0.1M β- Mercaptoethanol) as previously reported [7]. Tumour samples were prepared for lysis using a BioPulverizer and Cryo-cup grinder (BioSpec, USA) and lysed in RIPA buffer (Cell Signalling Technology, USA) supplemented with protease inhibitors (Mini, EDTA-free Protease Inhibitor Cocktail, Roche, UK) and phosphatase inhibitors (PhosSTOP, Roche, UK). Western blotting was performed as previously reported [7]. Antibodies used include AKT (pan), phospho AKT Serine 473, PARP, ß-actin or GAPDH, details noted in Supplementary Table 2. Detection was carried out using the ChemiDoc XRS+ (BioRad, UK). Densitometric analysis of band intensity of blots was carried out using Image J (NIH, USA).

**Supplementary Table 1 – Details of qPCR primers used in this study**

| Target | RNA type | Method | Manufacturer | Reference |
|---|---|---|---|---|
| **miR-187-3p** | MicroRNA | TaqMan | Applied Biosystems | 001193 |
| ***RNU48*** | MicroRNA | TaqMan | Applied Biosystems | 001006 |
| ***PTEN*** | mRNA | SYBR | Qiagen | QT00086933 |
| ***KDM5B*** | mRNA | SYBR | Qiagen | QT00060648 |
| ***DNMT1*** | mRNA | SYBR | Qiagen | QT00034335 |
| ***EP300*** | mRNA | SYBR | Qiagen | QT00094500 |
| ***B2M*** | mRNA | SYBR | Qiagen | QT00088935 |

**Supplementary Table 2 – Details Antibodies used**

| Target | Manufacturer | Reference | Dilution | Origin Species |
|---|---|---|---|---|
| **AKT (pan)** | Cell signalling | 4691 | 1:1000 | Rabbit mAb |
| **Phospho AKT** | Cell signalling | 4060 | 1:2000 | Rabbit mAb |
| **LC3b** | Cell signalling | 2775 | 1:1000 | Rabbit pAb |
| **PARP** | Cell signalling | 9542 | 1:1000 | Rabbit pAb |
| **PTEN** | Cell signalling | 9188 | 1:1000 | Rabbit mAb |
| **ß-actin** | Santa Cruz | SC-69879 | 1:10,000 | Mouse mAb |
| **GAPDH** | Ambion | AM4300 | 1:10,000 | Mouse mAb |
| **Mouse HRP (2$^{o}$)** | Dako | P0449 | 1:2000 | Rabbit pAb |
| **Rabbit HRP (2$^{o}$)** | Dako | P0448 | 1:2000 | Goat pAb |

*mAb*, Monoclonal antibody*; pAb,* Polyclonal antibody

**Supplementary Table *3.* Genes encoding PTEN transcription factors with miR-187 binding site**

| ***PTEN* transcription factor (TF)** | **MiR-187 seedposition on TF** | **Conservation of miR-187 binding site on TF in primates (%)** | **Known TF function in transcriptional regulation of *PTEN*** | **Reference** |
|---|---|---|---|---|
| *MAZ* | chr16:29819949 | 89 | Mechanism of PTEN regulation unknown | N/A |
| *NFIC* | chr19:3435127 | 89 | Mechanism of PTEN regulation unknown | N/A |
| *E2F1* | chr20:32264211 | 78 | One study demonstrated PTEN activation via binding to 5’ UTR | [8] |
| ***EP300*** | **chr22:41572511** | **78** | **Negatively regulates PTEN** | **[9]** |
| *BLC3* | chr19:45259490 | 67 | Mechanism of PTEN regulation unknown | N/A |
| *MYCBP2* | chr13:77835389 | 67 | Mechanism of PTEN regulation unknown | N/A |
| *NFIC* | chr19:3433521 | 67 | Mechanism of PTEN regulation unknown | N/A |
| *PHF8* | chrX:54069203 | 67 | Mechanism of PTEN regulation unknown | N/A |
| *PPARG* | chr4:23816000 | 67 | Positive regulator | [10] |
| ***KDM5B*** | **chr1:202698500** | **56** | **Histone demethylase which transcriptionally represses PTEN** | **[11]** |
| *MAZ* | chr16:29821826 | 56 | Mechanism of PTEN regulation unknown | N/A |
| *SRFBP1* | chr5:121356312 | 56 | Mechanism of PTEN regulation unknown | N/A |
| *PPARG* | chr4:23822068 | 56 | Positive regulator | [10] |
| *FOXP2* | chr7:113726400 | 44 | Mechanism of PTEN regulation unknown | N/A |
| *MAFK* | chr7:1580362 | 44 | Mechanism of PTEN regulation unknown | N/A |
| *WRNIP1* | chr6:2766547 | 44 | Mechanism of PTEN regulation unknown | N/A |
| ***DNMT1*** | **chr19:10248642** | **44** | **Negative regulator** | **[10]** |

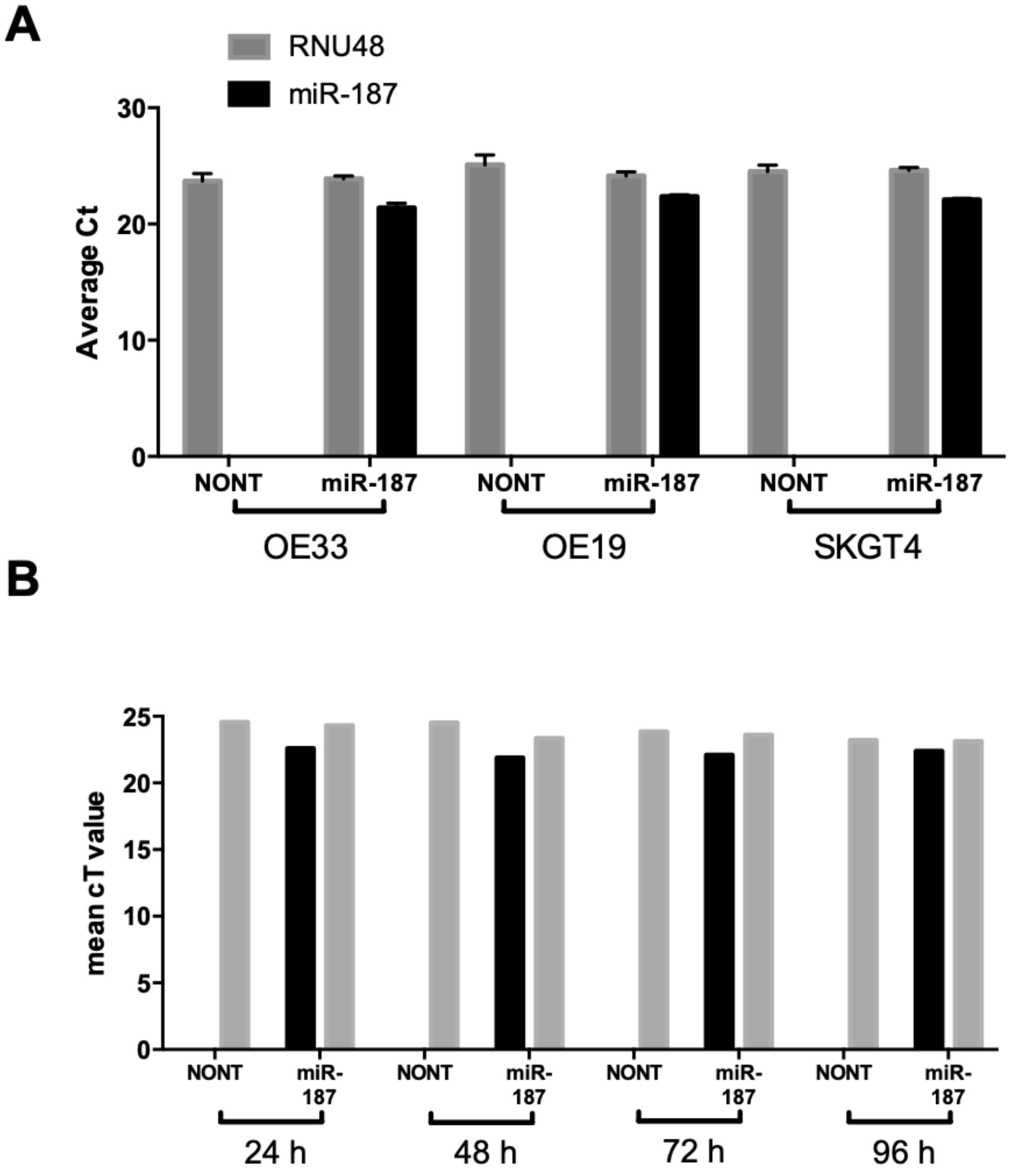


**Supplementary Figure 1 – Validation of mir-187 overexpression**

Cells were seeded into 6cm dishes and transiently transfected with 5 nM pre-miR-187. miR-187 expression levels were assessed 24 h post transfection for all cell lines (A) or for a series of timepoints post transfection for OE33 cell line (B) using TaqMan qPCR. MiR-187 expression was also assessed in mock transfected (NONT) control cells. The average Ct values for miR-187 and the endogenous control RNU48 are presented for each sample. Data represent the average of n=3 experimental repeats (A) or n=2 experimental repeats (B). Error bars represent mean ± SEM.

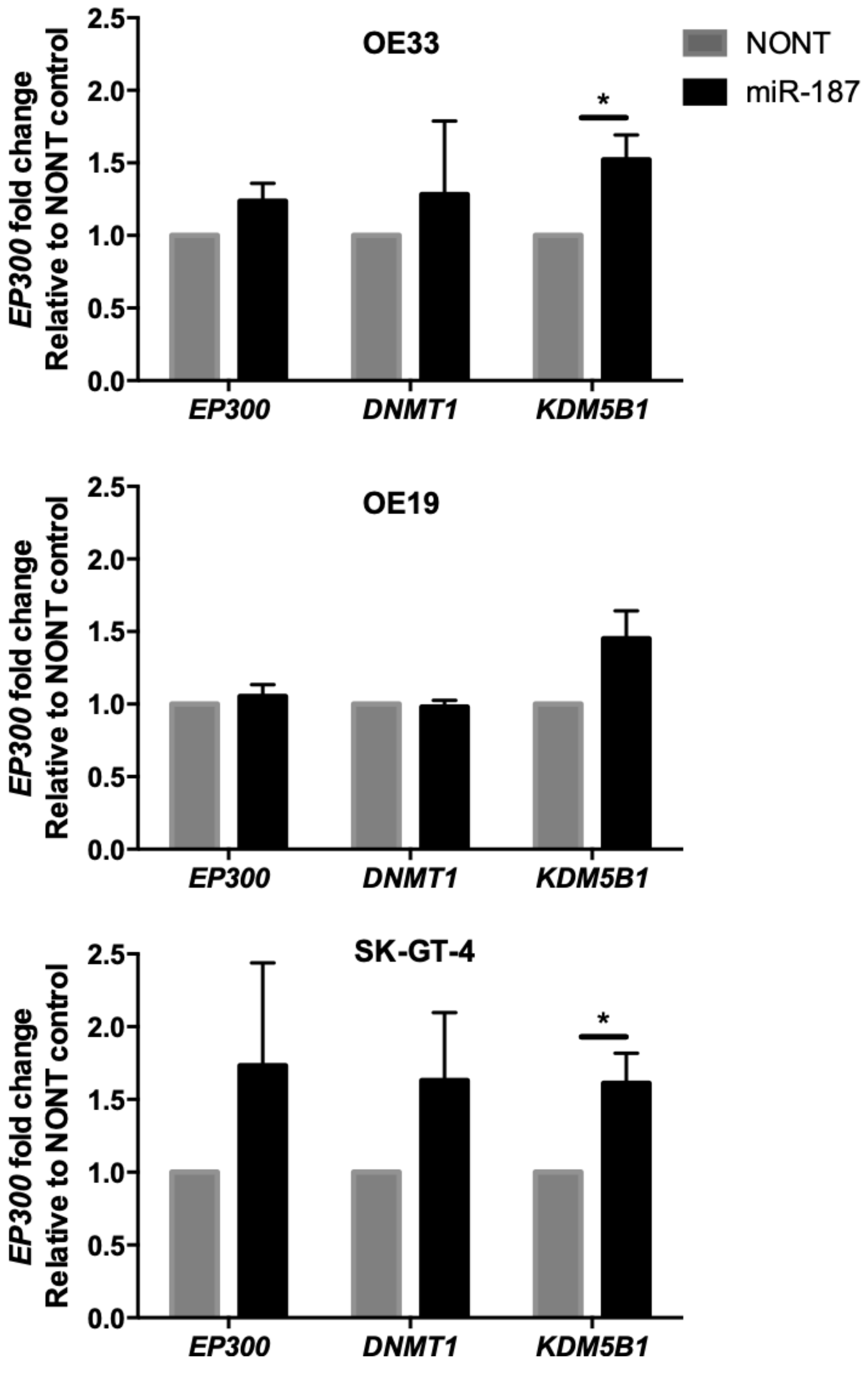


**Supplementary Figure 2 – Evaluation of potential transcriptional regulators of PTEN downstream of miR-187**

Cells were reverse transfected with 5 nM miR-187 precursor or mock transfected (Mock) and harvested 24 h post transfection. *EP300*, *DNMT1*, and *KDM5B* expression was analysed by qPCR Expression levels were normalised to mock transfected (Mock) control cells. *B2M* was used as a housekeeping gene. Histogram represents an average of n=3 independent experimental repeats. Error bars represent mean ± SEM. Statistical analysis was performed using the Student's t-test.

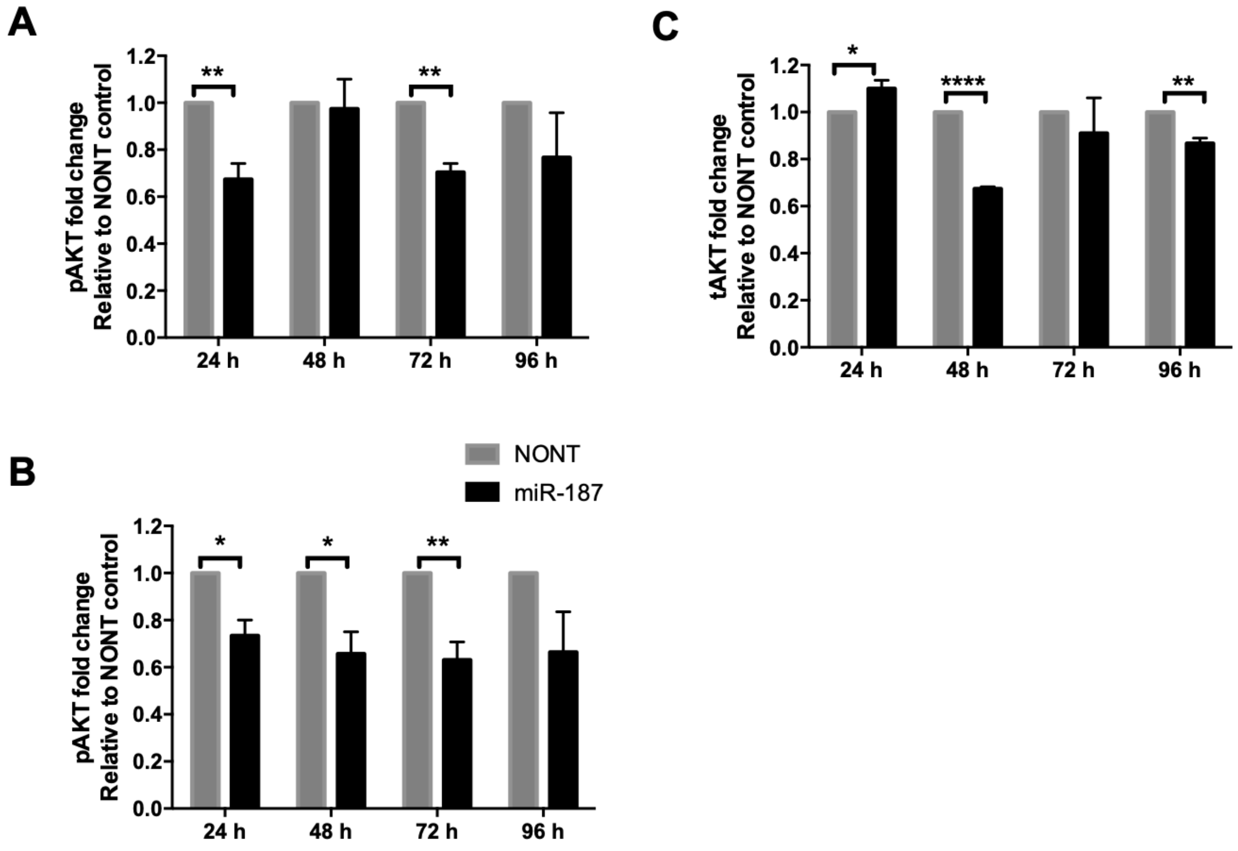


**Supplementary Figure 3 – Densitometry quantification of pAKT and tAKT**

Densitometry of western blots shown in Figure 2B. Band intensity was quantified using ImageJ. Histograms represent the average band intensity for pAKT normalised to tAKT (A) pAKT normalised to ß-actin control (B) and tAKT normalised to ß-actin control (C) All calculations are relative to the mock transfected control (Mock) for each time point. Histogram represents an average of n=3 independent experimental repeats. Error bars represent mean ± SEM. Statistical analysis was performed using the Student's t-test: $*p < 0.05$, $**p < 0.01$, $****p < 0.0001$.

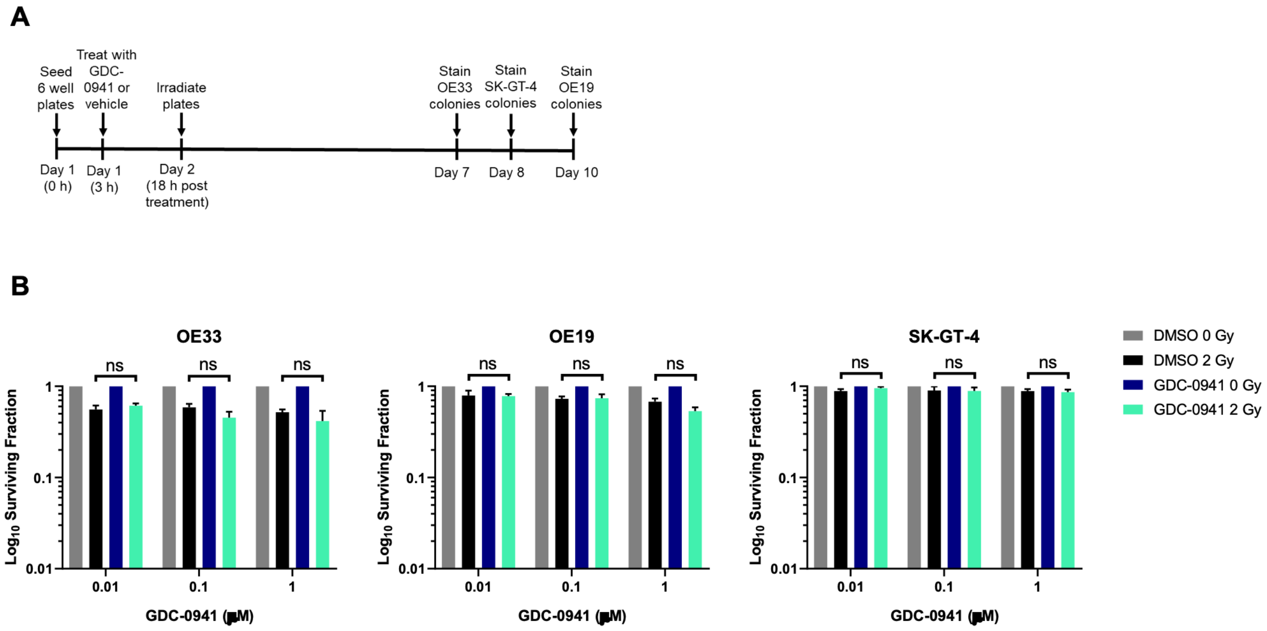


**Supplementary Figure 4 – GDC-0941 significantly reduced SK-GT-4 clonogenic survival as a single agent and has an additive effect in combination with physiologically relevant levels of irradiation**

Cells were seeded into 6 well plates for the clonogenic assay and treated with GDC-0941 or the corresponding vehicle control once adhered. Plates were transported to the irradiator and exposed to 0 Gy or 2 Gy irradiation doses 18 h post treatment. Colonies were stained and counted. The surviving fraction was calculated relative to the mock transfected (Mock) 0 Gy control. Treatment with GDC-0941 or vehicle control (DMSO) was removed at the time of staining. B Treatment with GDC-0941 or vehicle control (DMSO) was removed 24 h post irradiation and replaced with complete medium. (A) Treatment scheduling. (B) Survival fractions for all conditions, relative to GDC-0941-treated samples. Error bars represent mean ± SEM. Statistical analysis was performed using the Student's t-test.
